# Genetic Diversity Analysis of a Multi-Gene Bank Collection of Eggplant (*Solanum melongena* L.): Insights from Morphological Traits and SSR Markers

**DOI:** 10.1101/2024.04.01.587642

**Authors:** Patrick Louie G. Lipio, Ian Bien M. Oloc-Oloc, Timothy D. Guinto, Bryan Anthony A. Diamante, John Albert M. Caraan, Richard T. Hermoso, Visitacion C. Huelgas, Desiree M. Hautea

## Abstract

Multiple pests and diseases significantly affect eggplant (*Solanum melongena* L.) yield. Its principal pest, the eggplant fruit and shoot borer [EFSB, *Leucinodes orbonalis* Guenée (Lepidoptera: Crambidae)], damages plants from the early vegetative stage to the late fruiting stage, while infestation by the green leafhopper [*Amrasca biguttula* Ishida (Hemiptera: Cicadellidae)] stunts vegetative growth. Analyzing the diversity of the worldwide collections in different gene banks would allow for the effective use of stockpiled genetic resources for the breeding of insect-resistant eggplant. The genetic diversity and population structure of 96 accessions from four (4) germplasm collections were assessed using marker data from 20 poly-morphic microsatellite (SSR, simple sequence repeats) markers distributed among all 14 linkage groups. Phenotypes that were given focus included those linked to possible physical barriers to herbivory and possible presence of defensive compounds. The average number of alleles (n) across loci was 4.8, with an average polymorphism information content (PIC) value of 0.59, suggesting that the SSR loci were moderately informative. Population structure analysis yielded two subgroups with moderate to highly significant differentiation within each cluster (Fst_1_ and Fst_2_ = 0.23 and 0.14, respectively). Taken together, hierarchical clustering and population structure analysis were able to discriminate accessions by vegetative characters but were not able to effectively discriminate based on geo-graphic provenance, fruit phenotype, nor mean EFSB incidence.

## INTRODUCTION

Eggplant, *Solanum melongena* L., is a tropical perennial crop that is commercially cultivated as an annual crop in tropical and sub-tropical regions of the world. Considered the second most important solanaceous vegetable crop, producing up to 55.2 metric tons in 1.8M hectares, worldwide (FAOSTAT, 2021). In the Philippines, as of 2020, 242,730.40 metric tons have been produced in 21,780.33 hectares, making it the most produced vegetable in the country, followed closely by tomato (Philippine Statistics Authority, 2021).

Eggplant was largely proposed to have originated in Africa, where it later dispersed into the Middle East and then into Asia, where they were domesticated (Weese and Bohs, 2010). The domestication of eggplant was first proposed to be the around the Indio-Burma region (Vavilov, 1951), with later theories placing the area of domestication either in China or India (Daunay and Janick, 2007; Wang et al., 2008). Recent studies have found cases for both multiple domestication events in India, South China, and Southeast Asia (Meyer et al., 2012) and a single domestication event in Asia (Page et al., 2019). In either case, by the 8th century, domesticated eggplants spread eastward to Japan and westward to Africa and Europe (Prohens et al., 2005).

Previous studies on the global diversity of eggplants collected in genebanks revealed major clusters in India, Turkey, Southeast Asia, and Spain (Taher et al., 2017). Further diversity studies were also conducted among different eggplant populations, some of which included wild relatives (Naujeer, 2009; Demir et al., 2010; Sunseril et al., 2010; Munoz-Falcon et al., 2011; Begum et al., 2013; Ge et al., 2013; Naujeer, 2009; Caguiat and Hautea, 2014; Vilanova et al., 2014; Gramazio et al., 2019) and among major geographic clusters (Hurtado et al., 2012; Cericola et al., 2013). Diversity studies have also been conducted as means of screening for agronomically and commercially beneficial traits, such as phenolic content (Hanson et al., 2006; Prohens et al., 2007; Plazas et al., 2013; San Jose et al., 2013), heat tolerance (Uddin et al., 2014), and pest and disease resistance (Babu et al., 1999; Behera et al., 1999; Naegele et al., 2014; Asad et al., 2015).

Multiple pests and diseases significantly affect the crop’s yield, one of which is infestation with the eggplant fruit and shoot borer [EFSB, *Leucinodes orbonalis* Guenée (Lepidoptera: Crambidae)]. In the Philippines, high pest pressures have been recorded to reduce the yield of eggplant by up to 80%. EFSB affects the plant from the early vegetative stage to the late fruiting stage by damaging the shoots, hindering plant growth; damaging the flowers, preventing fruit formation; and damaging the fruits, rendering them unfit for human consumption (Francisco, 2014).

Integrating molecular and morphological data, the study aimed to assess the diversity and population structure within a collection of eggplant accessions sourced from different genebanks, laboratory holdings, and commercial seed producers in the Philippines. The accession panel consisted of morphologically diverse landraces, commercial varieties, open-pollinated entries, and crop wild relatives (CWRs) with varying geographical origins. Genotyping was performed through microsatellites (or SSRs) which have been instrumental in analyzing diversity among crop germplasm panels and selection for marker-assisted breeding. The findings of the study would be of great use in the development of outstanding eggplant breeding lines from available plant genetic resources. This would also be an important precedent for association studies geared towards finding genetic loci for resistance to EFSB.

## MATERIALS AND METHODS

### Plant materials

The seeds used in the experiment were sourced from the laboratory’s seed collection, genebanks, both local and foreign, and a commercial company. Genebanks include the National Plant Genetic Resources Laboratory (NPGRL), the United States Department of Agriculture (USDA), and Japan’s National Agriculture and Food Research Organization (NARO), while the commercial source was the East-West Seed Company. The morphological diversity observed among the eggplant accessions is illustrated in Figure 1, with the list of the accessions/germplasm and their respective sources found in Appendix A. Majority of the accessions selected for the study were done so based on physiological characteristics that may aid in the preventing herbivory, such as the presence of pigmented secondary metabolites and trichomes.

**Figure 1.**
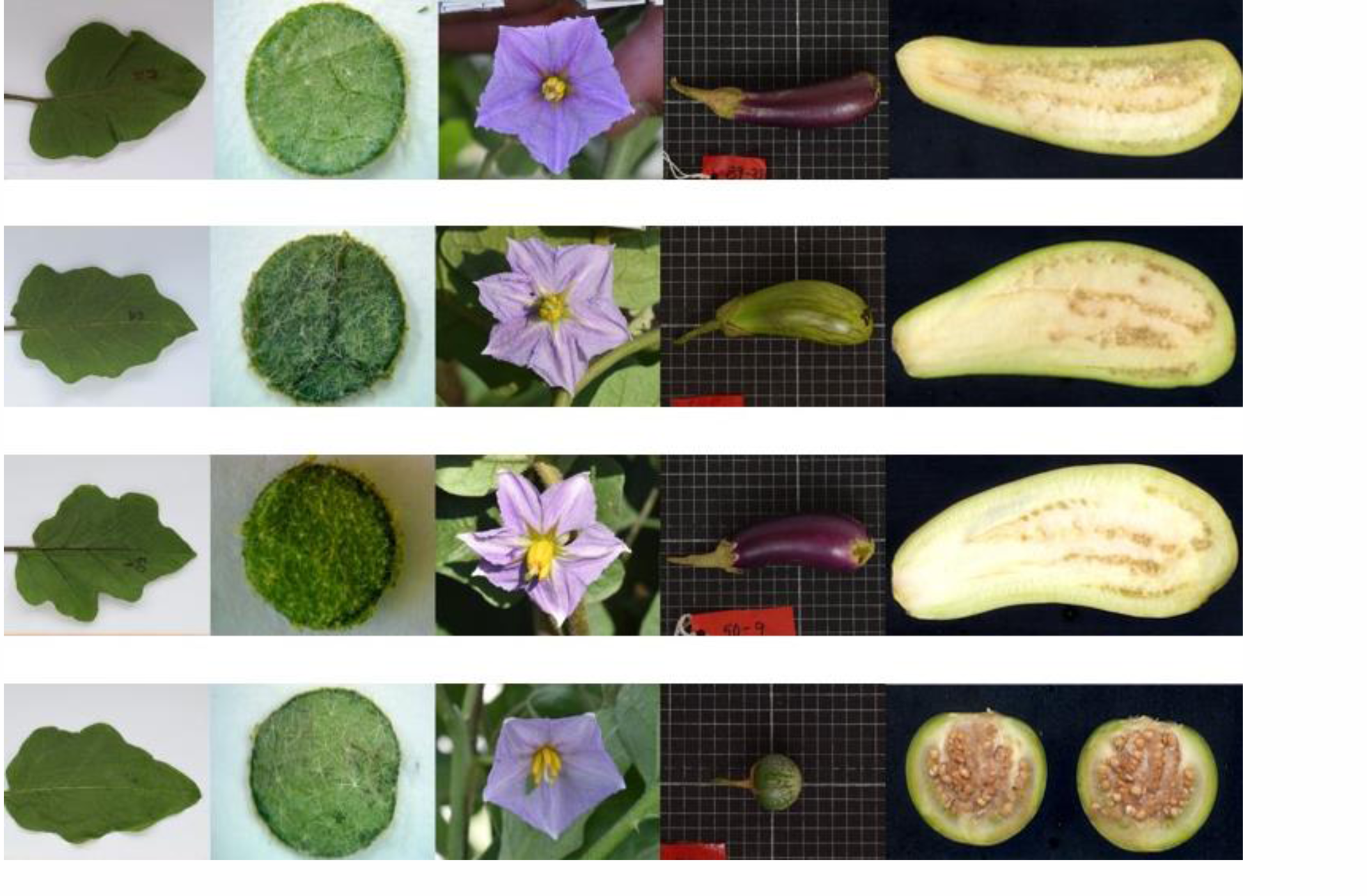
Morphological diversity of planting materials sourced from multiple gene banks and eggplant collections; (left to right) leaf, leaf trichomes, flower, immature fruit, and cross-section; (top to bottom) improved variety – Mara Mauve, landrace – GB 59477, unclassified – PI 362727, wild relative – PHL 9392.

### Crop Establishment, Experimental Design and Cultivation

Sowing for the 2018 – 2019 Dry Season Planting was conducted between 7 November 2018 and 24 November 2018 at the IPB BL2-2 greenhouse. Seedlings were transplanted on the 21 December 2018, for the first planting, and 4 January 2019, for the second planting, at Plot F (confined field area) of the IPB Central Experiment Station. For the 2018-2019 Dry Season Planting, all accessions were replicated twice, planted in rows following an alpha lattice design, separated into two large planting blocks, with each row composed of 15 plants of the same accession. Sowing for the 2019 – 2020 Dry Season Planting was conducted between 10 September 2019 and 28 October 2019 at the IPB BL2-2 greenhouse. Seedlings were transplanted on the 17 October 2019 for the first planting block and 28 October for the second planting block at Plot F (confined field area) of the IPB Central Experiment Station. For the 2019-2020 Dry Season Planting, all accessions were replicated twice, planted in rows following an alpha lattice design, with each row composed of 15 plants of the same accession.

### Phenotype Data Collection

After transplanting in the field, the vegetative characteristics, as stated by the IBPGR (1990), were measured once 50% of the plants in an accession on the field have started to flower. The specific vegetative characteristics possibly linked to insect resistance were reckoned, such as stem pubescence, leaf edge color and density of leaf pubescence. Fruit characteristics such as skin color at commercial ripeness, fruit flesh density, fruit calyx creasing, relative fruit calyx length, fruit pedicel length, fruit pedicel thickness and fruit calyx length were measured. The following morphological traits that were measured on a scale are specified as follows: Vegetative [Leaf Edge Color (Anthocyanin Presence) 0-9 (0-Absent, 9-Very Strong); Leaf Hair Density 0-9 (0-Glaborous, 9-Very Dense); Stem Pubescence 0-7 (0-Glaborous, 3-Weak, 5-Medium, 7-Strong)] and Reproductive [Fruit Pedicel Length (cm); Fruit Pedicel Thickness (cm); Fruit Calyx Length (cm); Fruit Curved Height (cm); Fruit Color at Commercial Ripeness 1-9 (1-Green, 2-Milk White, 3-Deep Yellow, 4-Fire Red, 5-Scarlet Red, 6-Lilac Grey, 7-Purple, 8-Purple Black, 9-Black); Fruit Flesh Density 1-9 (1-Spongy, 9-Very Dense); Fruit Calyx Creasing 1-9 (1-Very Weak, 9-Very Strong); Relative Fruit Calyx Length 1-9 (1-Very Short(<10%), 9-Very Long(>10%))]. Average and highest percent infestation, for the plants in a row, among the accessions planted in the field were also collected weekly for a period between 7 to 18 weeks after transplanting. This was done by inspecting the leaves, stems and fruits of each plant in a row for insect damage and scoring whether the observed damage for a specific insect exceeds the threshold.

### Genotyping

Leaf sample collection was done by punching four to five (4-5) leaf discs from mature leaves of the seedlings using 2 mL microcentrifuge tubes. Collections were immediately placed over to five leaf discs were collected from each plant. The microcentrifuge tube with the collected leaf discs were kept in –80°C until extraction. DNA was extracted following a modified CTAB method (CIMMYT, 2005). The resulting DNA samples were temporarily stored in 4°C, until 5 ng/µL working stocks were made from these samples. Afterwards, these stock DNA samples were placed in a -20°C refrigerator for long term storage.

SSR markers were selected based on previously reported SSR markers in Caguiat and Hautea (2014). A total of 20 SSR markers covering all 14 linkage groups, under reference map EW2009 [34] were amplified in the bulk samples. For the PCR assay, a 10μL PCR re-action mix (5.3uL Sigma Water; 1.0 10xPCR Buffer; 0.3uL 50uM Magnesium Chloride; 0.2uL 2.5uM DNTPs; 1.0uL 2uM Forward Primer; 1.0uL 2uM Reverse Primer; 0.2uL 5U/uL Taq Pol (Invitrogen); 1.0uL 2.5ng/uL Bulked DNA) was prepared for each bulk sample by first preparing a master mix for 155 PCR reactions. The PCR profile and markers used are presented in Appendix B. After amplification of markers, the samples were resolved in 8% acrylamide gel, prepared according to the protocol in Appendix 4. Loaded samples in 4X blue juice (Invitrogen, Carlsbad, CA) were run with molecular weight standards Marker VIII (Roche) at 100V for 60 min in 1X TBE. The gel was stained in ethidium bromide (ETBR) for 5 minutes, and washed for 1 minute in distilled water, prior to image capture using a gel documentation system (Enduro^TM^ Gel Documentation System, Labnet).

### Population Structure and Diversity Analysis

Scoring of the microsatellite data was based on an arbitrary numerical system from 1 to *n*, where *n* is the maximum number of polymorphic alleles observed for every marker. For putatively null amplifications and missing data, -9 was used as input. The allele frequency analysis was computed using CERVUS 3.0.7 Software (Kalinowski et al., 2007). These data were imported into the STRUCTURE 2.3 software (Pritchard et al., 2000) to obtain a Bayesian-based population structure model associated to origin and to assess the accuracy of the a priori assignments. As previously implemented by Cericola et al. (2013) K-values (i.e., hypothetical number of populations) from 1 to 10 were tested with a burn-in period of 50,000 rounds from ten independent simulations with three iterations for each K-value tested. The number of Markov Chain Monte Carlo (MCMC) replications after burn-in was 100,000. The resulting data files were then submitted to the server STRUCTURE Harvester (http://taylor0.biology.ucla.edu/structureHarvester/) to obtain ΔK values, their standard deviations, and log conversions (Earl and vonHoldt, 2012).

From this, the Evanno method (Evanno et al., 2005) was used to determine the best and most informative estimated number of populations within the germplasm. Since not all assignments can be grounded on verified genebank data, the admixture model was used to account for any overlaps in origin and to see relationships among variants that exist within the same postulated domestication origins.

### Cluster Analysis

SSR marker information and previously selected phenotype data were analyzed in the R programming environment (R Core Team, 2021). Null values for this step were all assigned the value “0”. K-means Clustering, and Principal Component Analysis was done for both the SSR marker data and the phenotype data employing the “kmeans” function and “prcomp” function respectively for each analysis. The optimal number of clusters for K-means Clustering was obtained by using the “fviz_nbclust” function in the “factoextra” package (Alboukadel and Fabian, 2020). Hierarchical clustering was also conducted to create a dendrogram, using the SSR marker data with the “hclust” function used to conduct the analysis. “kmeans”, “prcomp” and “hclust” previously mentioned are found in the base R package while the packages “cluster” (Maechler et al., 2021), “factoextra” (Alboukadel and Fabian, 2020) and “ggplot2” (Wickham, 2009) were used for the visual presentation of the data.

## RESULTS

### SSR Marker Data

From the obtained SSR marker data compiled in **Table 1**, the 97 accessions have an average of 4.8 alleles with the highest number of alleles observed in markers, EM4_1, ecm001, EM117 and the lowest number of alleles in markers ecm032, ecm023 and ecm070. The average PIC of the markers obtained was 0.59, meaning that the set of 20 SSR markers used in the panel were moderately informative. The most informative marker was EM117 with the PIC value of 0.79, while the lowest was for EM206 with PIC value of 0.44. The average value for observed heterozygosity was 0.06, while the value for the expected heterozygosity was 0.64, with values ranging from 0.000 to 0.349 and 0.464 to 0.816, respectively. The highest heterozygosity observed (Hobs) was 0.349 in marker EM206. The value for the fixation index (F) that was observed in the panel was 0.8461. All the loci conform to Hardy-Weinberg equilibrium.

**Table 1.**
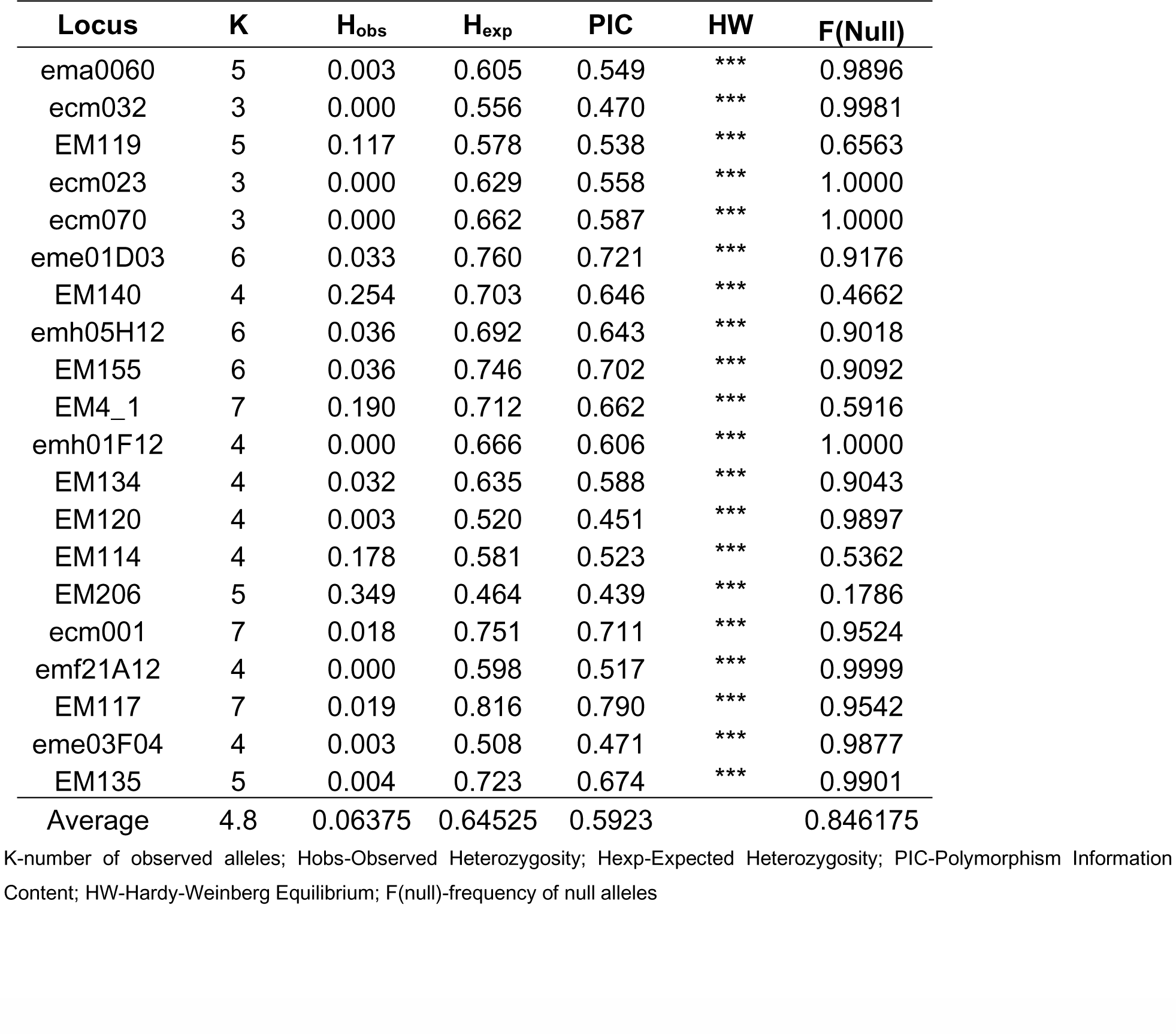
Summary of SSR Marker Data.

### Population Structure Analysis and K-means Clustering

Clustering using PCA with the SSR marker data was not able to yield any clearly de-fined separations in geographical groups, with many of the predefined clusters based on geographical origin overlapping with each other (**Figure 2**). Aside from this, clustering based on the classification of each accession was also not able to yield any defined clusters.

**Figure 2.**
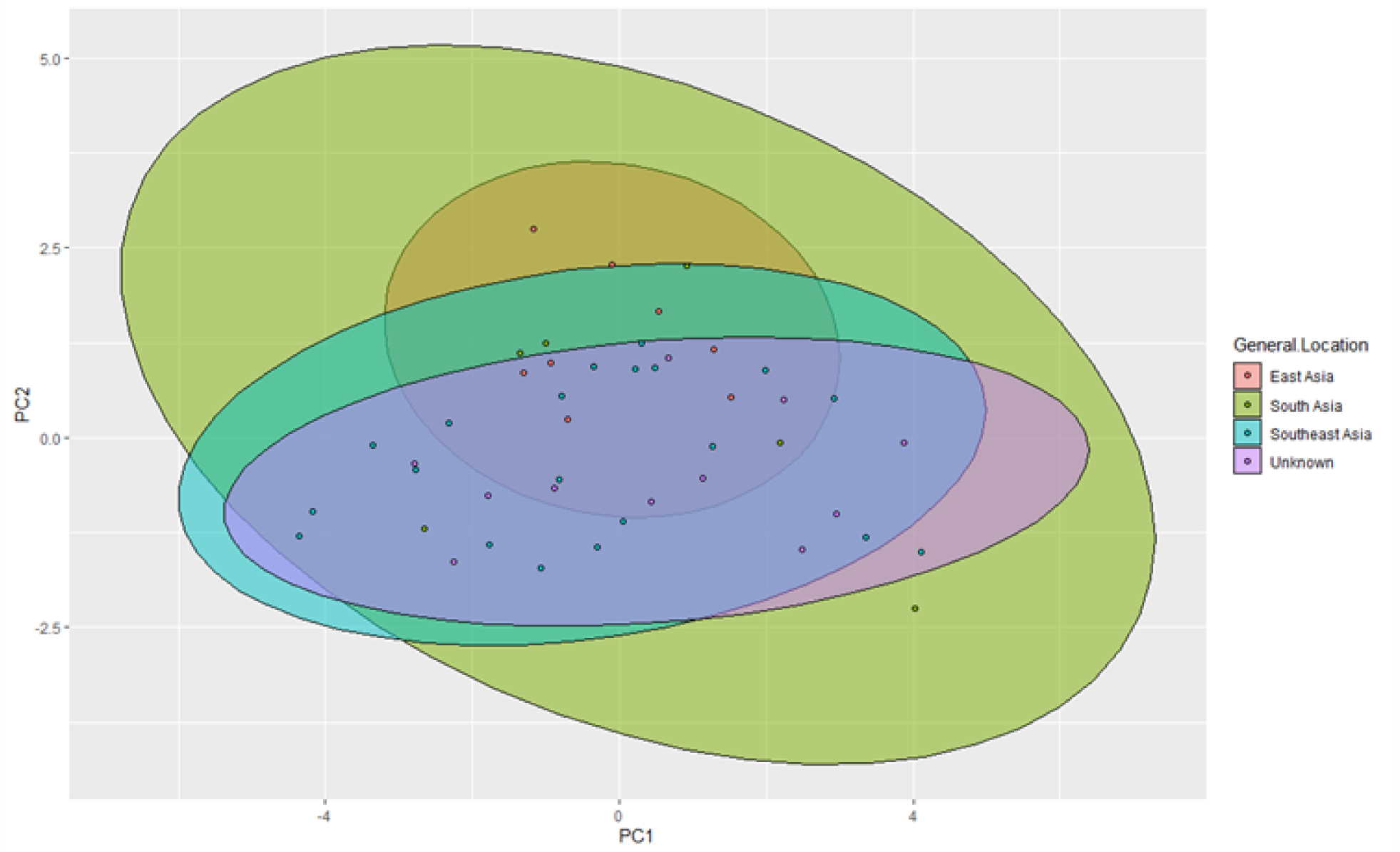
PCA clustering using SSR marker data based on geographic origin

The groupings through K-means clustering using the observed phenotypes have resulted in K=3 groups for both vegetative characteristics, and fruit characteristics and infestation incidence. These groups were based on the characteristics stated in the methodology and were separated based on whether they were vegetative characteristics or characteristics relating to the fruit.

Analysis of the population structure based on the phenotype data yielded more defined clusters. For vegetative data the two major defining components that differentiated the accessions were leaf edge color (38.7%) and leaf hair density (34.1%), both of which accounted for 72.8% of the variation found with regards to the observed vegetative data **(Figure 3)**. The most common score observed among the population for leaf hair density was 5 (Moderate Density), while the most common scores for stem pubescence and leaf edge color were 3 (Weak) and 5 (Moderate Leaf Edge Pigmentation), respectively.

**Figure 3.**
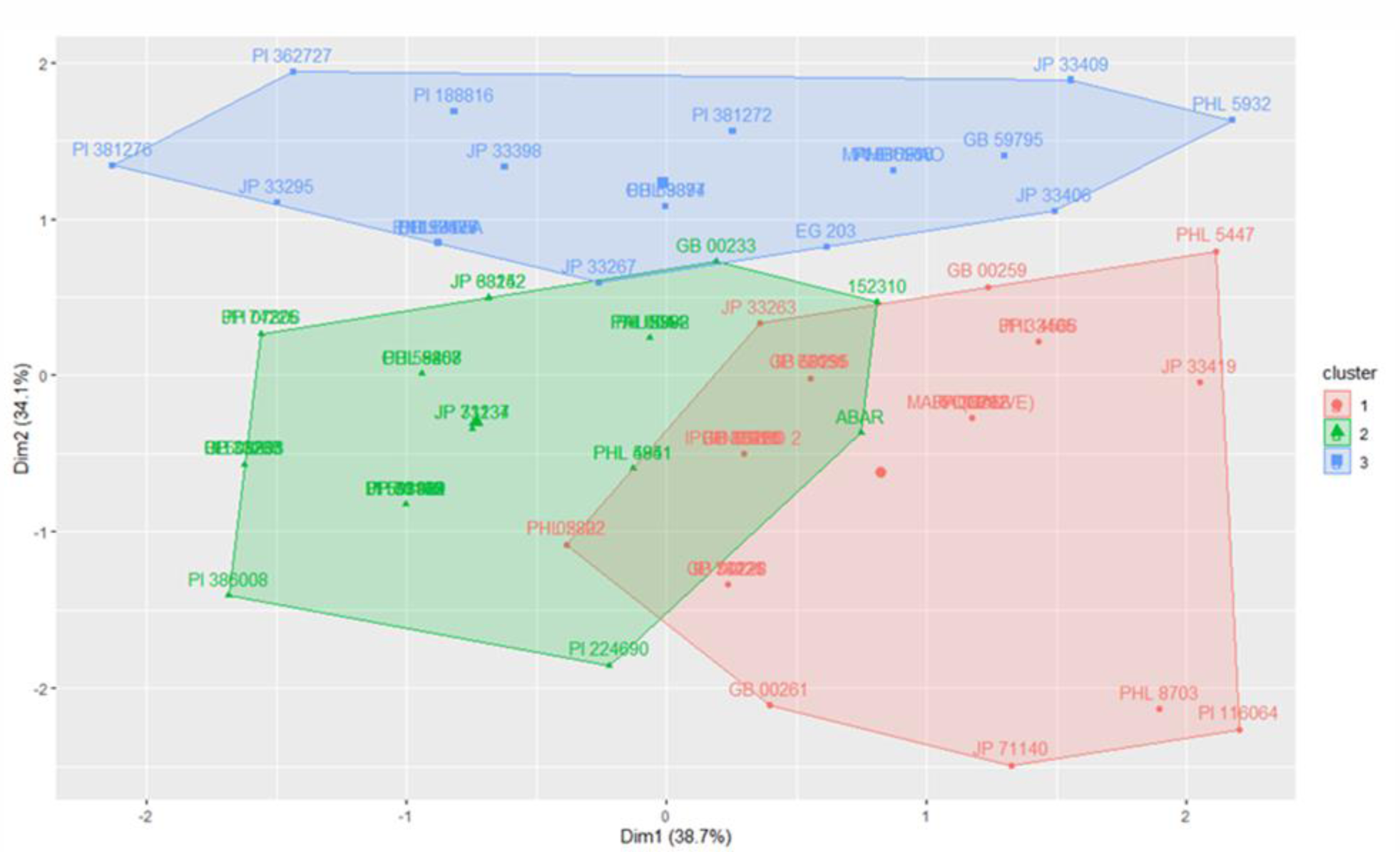
PCA Clustering using vegetative data with groupings based on K-means clustering

For the fruit data and EFSB fruit infestation data, the fruit calyx length (13.8%) and the maximum infestation occurrence (42%) were the two major defining components which accounted for 55.8% of the variation within the observed accessions (**Figure 4**). The most common score observed for fruit skin color at commercial ripeness was 8 (purple black) while the most common scores for fruit density, calyx creasing and relative fruit calyx length were 5 (Moderate Density), 5 (Moderate Creasing) and 3 (Moderately Short), respectively. For EFSB fruit infestation maximum, 100% infestation in all plants in a row was observed in the accessions GB59295 and PI362727, while JP33351 had the lowest observed infestation of plants in a row at 10%. For EFSB fruit infestation average, PHL4841 had the highest average infestation at 48.43%, among all plants throughout the entire observed period, while JP72036 had the lowest average infestation at 2.6%.

**Figure 4.**
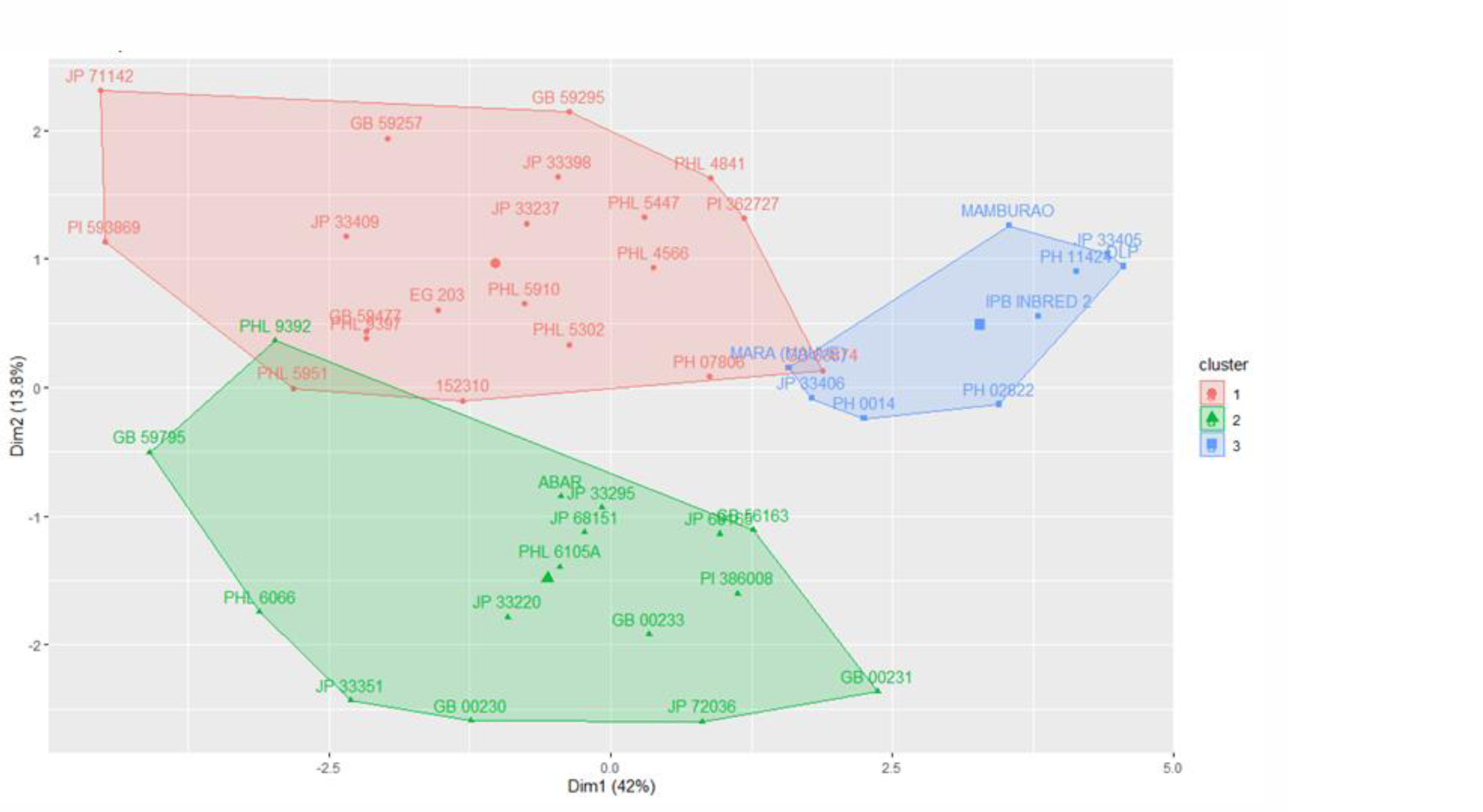
PCA clustering using fruit and EFSB infestation data with groupings based on K-means clustering

K-means clustering, and population structure analysis yielded two subgroups with moderate to highly significant differentiation within each cluster (Fst1 and Fst2 = 0.23 and 0.14, respectively). The result of K-means clustering using the SSR maker data has resulted in the K = 3 clusters (**Figure 5**). From both the STRUCTURE analysis and hierarchical clustering, it was noted that the accessions from the USDA did not cluster together, but with the accessions from other genebanks, while the accessions from the NARO genebank clustered together, but some accessions also grouped with accessions from other sources. Accessions sourced from Philippine collections tend to cluster together. Overall, there seems to be no clearly defined population structure among the accessions selected with some level of admixture among the groups defined by the dendrogram, with regards to the SSR marker data. With regards to the correlation between the accessions in the K subgroups based on the SSR markers obtained in STRUCTURE and the accessions in the K subgroups obtained from using the k-means function in R, there was a 56% correlation between the previous phenotype K subgroup 1 and genotype K subgroup 1, while the correlation between the phenotype K subgroup 2 and genotype K subgroup 2 was at 87%. This contrasts with the third subgroup, where the correlation between the genotype and phenotype data was only at 14%, with most of them being classified instead under the second genotype-based K subgroup.

**Figure 5.**
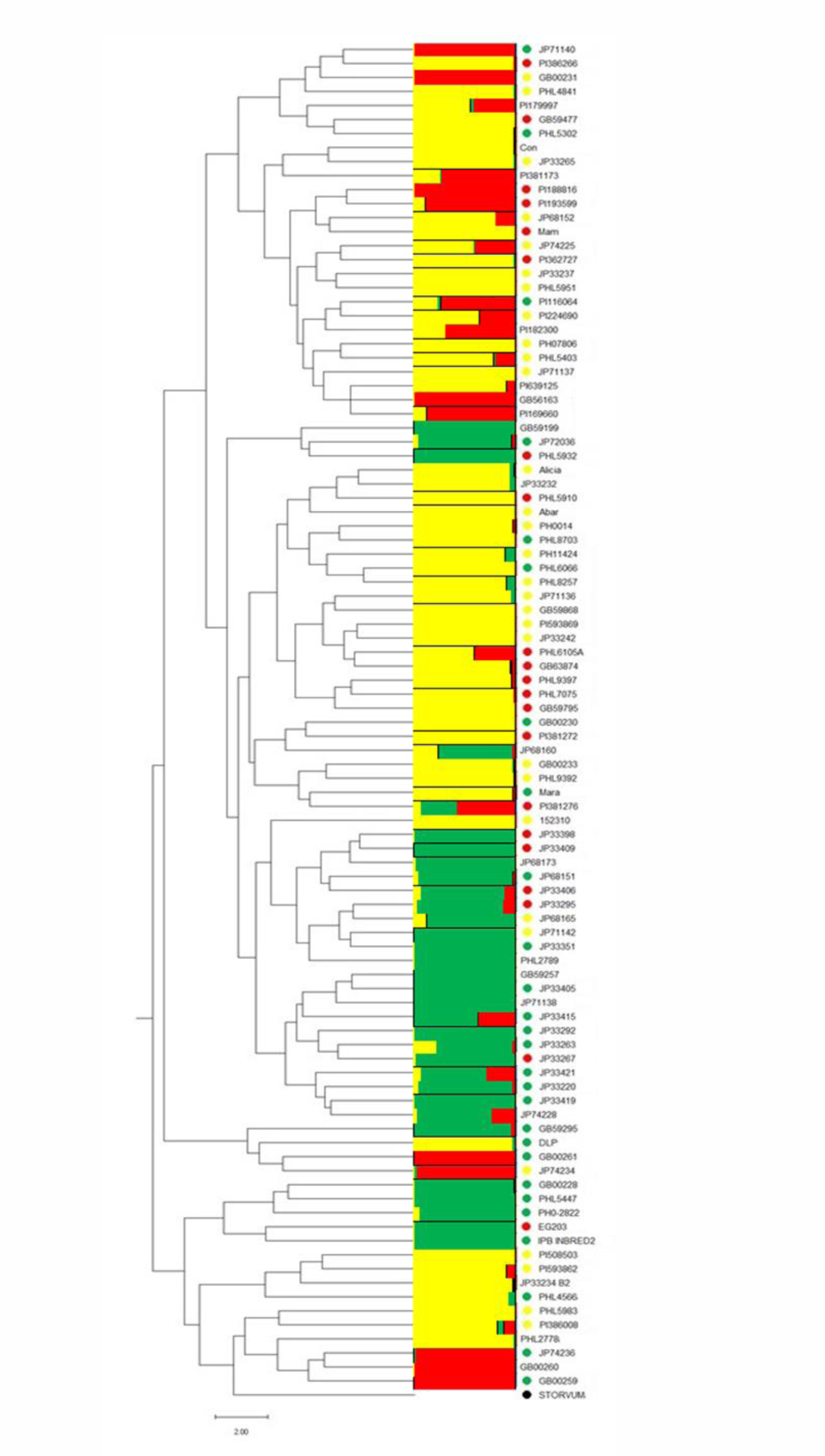
Dendrogram based on hierarchical clustering (K=3) of genetic marker data (bar plot), and obtained (K=3) PCA clustering based on vegetative characters (right points).

## DISCUSSION

The average number of alleles (n) obtained in the study (n = 4.8) was found to be higher than the previous study dealing with Philippine cultivars (n = 2.27) (Caguiat and Hautea, 2014), though lower than those conducted using a much larger and more diverse population, such as Cericola et al. (n = 5.8) (2013) and Hurtado et al. (n = 9.1) (2012). The value is also comparable to a previous study by Vilanova et al. (n = 4.3) (2014) which dealt with the diversity of landraces in Spain, a secondary center of diversity, Demir (n = 4.8) (2010), which utilized Turkish accessions, and Gramazio et al. (n = 4.6) (2019), who studied the diversity among peninsular and insular Greek cultivars. The assumption would be, that were it not for the addition of foreign accessions from various collections, local and foreign, the average number of alleles would hold closer to Caguiat and Hautea (2014), whose study dealt with Philippine accessions and wild relatives and Ge et al. (n = 3.1) (2013), whose study dealt exclusively with locally collected Chinese accessions. Though the average number of alleles for the study could be higher, were the accessions, obtained from the genebank collections, not limited to those with known resistance to EFSB, or those with vegetative and fruit characteristics that may serve as a barrier to insect herbivory. Another explanation for the low average would be the self-fertilizing nature of the species (Pessarakli and Driss, 2004). This highlights the benefit of using genebank collections and the careful assessment of the effective panel size and coverage of diversity in finding accessions that are excellent candidates for breeding programs (Taher et al., 2017).

PIC values for the set of 20 SSR markers used were found to be moderately informative (PIC = 0.59). Compared to previous studies which have lower PIC scores (Demir et al., 2010), where only 56% of the markers reached the slightly informative level, the moderately informative PIC scores have more in common with other studies that have selected their markers based on previous studies (Demir et al., 2010; Gramazio et al., 2019).

The fixation index (Fst) obtained is comparable to other diversity studies Vilanova et al. (2014) (Fst = 0.88), Cericola et al. (2013) (Fst = 0.86) and Gramazio et al (2019) (Fst = 0.79). This shows that eggplant to be an outcrossing species, though previous studies have stated the autogamous nature of eggplant as means of explaining the low average number of alleles per marker (Pessarakli and Driss, 2004). This may show the significance of factors, such as pollinators (Sambandam, 1964), as well as the floral morphology play in the reproduction of eggplant (Davidar et al., 2015). The distinct difference between the observed heterozygosity (Hobs = 0.06) and expected heterozygosity (Hexp = 0.64) is expected due to the autogamous nature of the plant (Pessarakli and Driss, 2004) as well as a possible narrow genetic base caused by inbreeding (Ali et al., 2011; Nunome et al., 2003).

As in previous studies (Cericola et al., 2013: Hurtado et al., 2012), there is a weak correlation between the genetic marker data and the phenotypic data, with regards to studies dealing with diverse populations, as compared to more specific and specialized populations (Munoz-Falcon et al., 2011; Prohens et al., 2005; Vilanova et al., 2014). This can be observed with the inability of the data in STRUCTURE analysis to correlate with the clustering obtained with the phenotype data, for the cases of fruit data and EFSB infestation data. STRUCTURE analysis was also unable to separate the population based on geographic provenance, unlike Cericola et al. (2013).

## CONCLUSION

From the study, K-means and STRUCTURE analysis were able to discriminate accessions by leaf pigmentation but were not able to effectively discriminate based on geographic provenance, fruit phenotype, nor mean EFSB incidence. No significantly defined clusters based on the latter three parameters were observed, which mutually suggest population admixture and the insufficiency of the marker data to differentiate based on traits largely believed to be polygenic. Genetic indices such as PIC, Fst, and number of observed alleles were variably comparable to previous population structure studies done in eggplant. Factors such as germplasm panel size and marker density (i.e., number of markers genotyped and chromosomal distribution) might have been critical differentiating determinants of success. Previous observations with regards to eggplant population dynamics held consistently with the experimental germplasm i.e., weak correlation between eggplant genotype and phenotype data as germplasm diversity increases, and that despite having notable capacity to outcross, eggplant physiologically prefers to self-fertilize.

## ACKNOWLEDGMENTS

The authors would like to recognize the contributions of Ms. Marilyn de Vera in the conduct of DNA extraction and SSR marker analysis. This study was funded by the Department of Science and Technology Philippine Council for Agriculture Aquatic and Natural Resources Research and Development Grants in Aid (DOST-PCAARRD-GIA) with matching funds by UPLB. This project was implemented under UPLB Project Code N91832.

