## Supplemental Tables for "Genetic Diversity Analysis of a Multi-Gene Bank Collection of Eggplant (*Solanum melongena* L.): Insights from Morphological Traits and SSR Markers"

**SUPPLEMENTAL TABLES AND FIGURES**

**Table A1**. List of Eggplant and Wild Relative Accessions from Various Genebank Collections

|  | Accession | Source | Geographic Origin | Taxon | Classification |
| --- | --- | --- | --- | --- | --- |
| 1 | 152310 | Vegetables Collection | Philippines | *Solanum melongena L.* | Breeding Line |
| 2 | Abar | Genetics Collection | Philippines | *Solanum melongena L.* | Open Pollinated Variety |
| 3 | Alicia | Genetics Collection | Philippines | *Solanum melongena L.* | Commercial Variety |
| 4 | Con | Genetics Collection | Philippines | *Solanum melongena L.* | Open Pollinated Variety |
| 5 | DLP | Genetics Collection | Philippines | *Solanum melongena L.* | Modern Cultivated Variety |
| 6 | EG203 | Genetics Collection | Sri Lanka | *Solanum melongena L.* | Breeding Line |
| 7 | GB00228 | East West Seed Company | Unknown | *Solanum melongena L.* | Unknown |
| 8 | GB00230 | East West Seed Company | Unknown | *Solanum melongena L.* | Unknown |
| 9 | GB00231 | East West Seed Company | Unknown | *Solanum melongena L.* | Unknown |
| 10 | GB00233 | East West Seed Company | Unknown | *Solanum melongena L.* | Unknown |
| 11 | GB00259 | East West Seed Company | Unknown | *Solanum aethiopicum L.* | Wild Relative |
| 12 | GB00260 | East West Seed Company | Unknown | *Solanum aethiopicum L.* | Wild Relative |
| 13 | GB00261 | East West Seed Company | Unknown | *Solanum aethiopicum L.* | Wild Relative |
| 14 | GB56163 | NPGRL Genebank | Unknown | Unknown | Unknown |
| 15 | GB59199 | NPGRL Genebank | Philippines | *Solanum melongena L.* | Unknown |
| 16 | GB59257 | NPGRL Genebank | Unknown | Unknown | Unknown |
| 17 | GB59295 | NPGRL Genebank | Philippines | *Solanum melongena L.* | Landrace |
| 18 | GB59477 | NPGRL Genebank | Philippines | *Solanum melongena L.* | Landrace |
| 19 | GB59795 | NPGRL Genebank | Philippines | *Solanum melongena L.* | Landrace |
| 20 | GB59868 | NPGRL Genebank | Philippines | *Solanum melongena L.* | Landrace |
| 21 | GB63874 | NPGRL Genebank | Philippines | *Solanum melongena L.* | Unknown |
| 22 | IPB INBRED2 | Genetics Collection | Philippines | *Solanum melongena L.* | Breeding Line |
| 23 | JP33220 | NARO Genebank | Japan | *Solanum melongena L.* | Breeding Line |
| 24 | JP33232 | NARO Genebank | Japan | *Solanum melongena L.* | Unknown |
| 25 | JP33234 | NARO Genebank | Japan | *Solanum melongena L.* | Unknown |
| 26 | JP33237 | NARO Genebank | Japan | *Solanum melongena L.* | Landrace |
| 27 | JP33242 | NARO Genebank | Japan | *Solanum melongena L.* | Landrace |
| 28 | JP33263 | NARO Genebank | Unknown | *Solanum melongena L.* | Unknown |
| 29 | JP33265 | NARO Genebank | Japan | *Solanum melongena L.* | Unknown |
| 30 | JP33267 | NARO Genebank | Japan | *Solanum melongena L.* | Unknown |
| 31 | JP33292 | NARO Genebank | Japan | *Solanum melongena L.* | Unknown |
| 32 | JP33295 | NARO Genebank | Japan | *Solanum melongena L.* | Unknown |
| 33 | JP33351 | NARO Genebank | Taiwan | *Solanum melongena L.* | Breeding Line |
| 34 | JP33398 | NARO Genebank | Bangladesh | *Solanum melongena L.* | Unknown |
| 35 | JP33405 | NARO Genebank | Bangladesh | *Solanum melongena L.* | Unknown |
| 36 | JP33406 | NARO Genebank | Bangladesh | *Solanum melongena L.* | Unknown |
| 37 | JP33409 | NARO Genebank | Bangladesh | *Solanum melongena L.* | Unknown |
| 38 | JP33415 | NARO Genebank | USSR | *Solanum melongena L.* | Unknown |
| 39 | JP33419 | NARO Genebank | Romania | *Solanum melongena L.* | Unknown |
| 40 | JP33421 | NARO Genebank | Romania | *Solanum melongena L.* | Unknown |
| 41 | JP68151 | NARO Genebank | Japan | *Solanum melongena L.* | Landrace |
| 42 | JP68152 | NARO Genebank | Japan | *Solanum melongena L.* | Landrace |
| 43 | JP68160 | NARO Genebank | Japan | *Solanum melongena L.* | Unknown |
| 44 | JP68165 | NARO Genebank | Japan | *Solanum melongena L.* | Landrace |
| 45 | JP68173 | NARO Genebank | Turkey | *Solanum melongena L.* | Unknown |
| 46 | JP71136 | NARO Genebank | Japan | *Solanum melongena L.* | Landrace |
| 47 | JP71137 | NARO Genebank | China | *Solanum melongena L.* | Unknown |
| 48 | JP71138 | NARO Genebank | France | *Solanum melongena L.* | Landrace |
| 49 | JP71140 | NARO Genebank | Unknown | *Solanum melongena L.* | Unknown |
| 50 | JP71142 | NARO Genebank | Thailand | *Solanum melongena L.* | Landrace |
| 51 | JP72036 | NARO Genebank | China | *Solanum melongena L.* | Landrace |
| 52 | JP74225 | NARO Genebank | Japan | *Solanum melongena L.* | Landrace |
| 53 | JP74228 | NARO Genebank | Unknown | *Solanum melongena L.* | Unknown |
| 54 | JP74234 | NARO Genebank | Japan | *Solanum melongena L.* | Unknown |
| 55 | JP74236 | NARO Genebank | Japan | *Solanum melongena L.* | Unknown |
| 56 | Mam | Genetics Collection | Philippines | *Solanum melongena L.* | Open Pollinated Variety |
| 57 | Mara | Genetics Collection | Philippines | *Solanum melongena L.* | Open Pollinated Variety |
| 58 | PH07806 | East West Seed Company | Unknown | *Solanum melongena L.* | Landrace |
| 59 | PH0014 | East West Seed Company | Unknown | *Solanum melongena L.* | Open Pollinated Variety |
| 60 | PH02822 | East West Seed Company | Unknown | *Solanum melongena L.* | Landrace |
| 61 | PH11424 | East West Seed Company | Unknown | *Solanum melongena L.* | Unknown |
| 62 | PHL5951 | NPGRL Genebank | Philippines | *Solanum melongena L.* | Unknown |
| 63 | PHL2778 | NPGRL Genebank | Philippines | *Solanum melongena L.* | Landrace |
| 64 | PHL2789 | NPGRL Genebank | Philippines | *Solanum melongena L.* | Landrace |
| 65 | PHL4566 | NPGRL Genebank | Philippines | *Solanum melongena L.* | Landrace |
| 66 | PHL4841 | NPGRL Genebank | Philippines | *Solanum melongena L.* | Unknown |
| 67 | PHL5302 | NPGRL Genebank | Philippines | *Solanum melongena L.* | Unknown |
| 68 | PHL5447 | NPGRL Genebank | Philippines | *Solanum melongena L.* | Unknown |
| 69 | PHL5403 | NPGRL Genebank | Philippines | *Solanum melongena L.* | Unknown |
| 70 | PHL5910 | NPGRL Genebank | Philippines | *Solanum melongena L.* | Unknown |
| 71 | PHL5932 | NPGRL Genebank | Philippines | *Solanum melongena L.* | Unknown |
| 72 | PHL5951 | NPGRL Genebank | Philippines | *Solanum melongena L.* | Unknown |
| 73 | PHL5983 | NPGRL Genebank | Philippines | *Solanum melongena L.* | Unknown |
| 74 | PHL6066 | NPGRL Genebank | Philippines | *Solanum melongena L.* | Unknown |
| 75 | PHL6105A | NPGRL Genebank | Philippines | *Solanum melongena L.* | Unknown |
| 76 | PHL7075 | NPGRL Genebank | Vietnam | *Solanum melongena L.* | Unknown |
| 77 | PHL8257 | NPGRL Genebank | Philippines | *Solanum melongena L.* | Unknown |
| 78 | PHL8703 | NPGRL Genebank | Philippines | *Solanum melongena L.* | Unknown |
| 79 | PHL9392 | NPGRL Genebank | Unknown | *Solanum linociera* | Wild Relative |
| 80 | PHL9397 | NPGRL Genebank | Unknown | *Solanum lasciocarpum Dunal* | Wild Relative |
| 81 | PI116064 | USDA Genebank | India | *Solanum melongena L.* | Unknown |
| 82 | PI169660 | USDA Genebank | Turkey | *Solanum melongena L.* | Unknown |
| 83 | PI179997 | USDA Genebank | India | *Solanum melongena L.* | Unknown |
| 84 | PI182300 | USDA Genebank | Turkey | *Solanum melongena L.* | Unknown |
| 85 | PI188816 | USDA Genebank | Philippines | *Solanum melongena L.* | Unknown |
| 86 | PI193599 | USDA Genebank | Ethiopia | *Solanum melongena L.* | Unknown |
| 87 | PI224690 | USDA Genebank | Myanmar | *Solanum melongena L.* | Unknown |
| 88 | PI362727 | USDA Genebank | India | *Solanum melongena L.* | Unknown |
| 89 | PI381173 | USDA Genebank | India | *Solanum melongena L.* | Unknown |
| 90 | PI381272 | USDA Genebank | India | *Solanum melongena L.* | Unknown |
| 91 | PI381276 | USDA Genebank | India | *Solanum melongena L.* | Unknown |
| 92 | PI386008 | USDA Genebank | Japan | *Solanum melongena L.* | Hybrid |
| 93 | PI386266 | USDA Genebank | India | *Solanum melongena L.* | Unknown |
| 94 | PI508503 | USDA Genebank | Korea | *Solanum melongena L.* | Cultivated Variety |
| 95 | PI593862 | USDA Genebank | Thailand | *Solanum melongena L.* | Unknown |
| 96 | PI593869 | USDA Genebank | Thailand | *Solanum melongena L.* | Unknown |
| 97 | PI639125 | USDA Genebank | United States | *Solanum melongena L.* | Unknown |

**Table B1. Microsatellite markers used for the Genotyping**

| **Marker name** | **Linkage group** | **Sequence** | **Annealing temperature (°C)** | **Expected band size, bp** |
| --- | --- | --- | --- | --- |
| eme01D03 | LG01 | F – 5’ aagaatcggtcctctttgcattgt- 3’ | 70 | 223-242 |
|  |  | R – 5’ tgcttttcacctctccgctatctc- 3’ |  |  |
| EM135 | LG01 | F – 5’ atcctgttgctgctcattttcctc- 3’ | 58 | 260 |
|  |  | R – 5’ aggaggatccaagaggtttgttga- 3’ |  |  |
| EM4_1 | LG02 | F – 5’ gcatacagcaaatcccagcaaata – 3’ | 63 | 172 |
|  |  | R – 5’ tgtgagtatgggattctggtcgtt – 3 |  |  |
| Ecm001 | LG03 | F – 5’cccttacgcaatttacacttcccc- 3’ | 65 | 227-229 |
|  |  | R – 5’atcaatggcgtcacctctctctct- 3’ |  |  |
| EM155 | LG03 | F – 5’caaaagataaaaagctgccggatg- 3’ | 65 | 248 |
|  |  | R – 5’catgcgtgagttttggagagagag- 3’ |  |  |
| Ema0060 | LG04 | F – 5’gctcggtatggcataagtttggag- 3’ | 66 | 393-397 |
|  |  | R – 5’gtttagcttccccattgtacccctgaac- 3’ |  |  |
| EM119 | LG04 | F – 5’ccccaccccatttgtgttatgtt- 3’ | 65 | 201 |
|  |  | R – 5’acccgagagctatggagtgttctg- 3’ |  |  |
| EM117 | LG05 | F – 5’gatcatcactggtttgggctacaa- 3’ | 65 | 123 |
|  |  | R – 5’aggggagaggaaacttgattggac- 3’ |  |  |
| Ecm070 | LG05 | F – 5’ctcaaaatccatggaggttttcca- 3’ | 68 | 226-230 |
|  |  | R – 5’tcctgttccgatttagctctcacc- 3’ |  |  |
| EM134 | LG06 | F – 5’ agtaagggaaagtgctgacgaagg – 3’ | 65 | 168 |
|  |  | R – 5’ cagagtcatcgttatggggaggtt – 3’ |  |  |
| Ecm023 | LG07 | F – 5’ tccaaacagtgtgacaaagcatga – 3’ | 69 | 253-257 |
|  |  | R – 5’ cccacttgggtatcactaagccac – 3’ |  |  |
| EM120 | LG08 | F – 5’ggatcaactgaagagctggtggtt- 3’ | 65 | 160 |
|  |  | R – 5’cagagcttcaatgttccatttcaca- 3’ |  |  |
| EM114 | LG08 | F – 5’agcctaaacttggttggtttttgc- 3’ | 65 | 221 |
|  |  | R – 5’gaagctttaagagccttctatgcag- 3’ |  |  |
| EM206 | LG09 | F – 5’atcttaatcttcctgctcctgttg- 3’ | 60 | 183 |
|  |  | R – 5’ctggaaacaagctcgctaccaaat- 3’ |  |  |
| Ecm032 | LG09 | F – 5’ tgcatgcaaaagagtgagcctaag – 3’ | 66 | 277-280 |
|  |  | R – 5’ ccctctctactgtgcccatgattt – 3’ |  |  |
| Emh01F12 | LG10 | F – 5’tcactcagccttattgccatgtgt- 3’ | 63 | 236 |
|  |  | R – 5’agtttgggaataggaaggagctgg- 3’ |  |  |
| EM140 | LG11 | F – 5’ccaaaacaatttccagtgactgtgc- 3’ | 65 | 268 |
|  |  | R – 5’gaccagaatgcccctcaaattaaa- 3’ |  |  |
| Eme03F04 | LG12 | F – 5’tatgacgacagacgtaaagcgacc- 3’ | 73 | 215-261 |
|  |  | R – 5’cagagttttgccatctgtgtcgag- 3’ |  |  |
| *Emf21A12 | LG13 | F – 5’ atcctggccatgtttctccattta – 3’ | 65 | 169-205 |
|  |  | R – 5’ cgtttgctttctaggagacttttagcc – 3’ |  |  |
| emh05H12 | LG14 | F – 5’ggtcactgctcttagtttctgcaa- 3’ | 68 | 149-203 |
|  |  | R – 5’cagagcagcgatcctttcttcatt- 3’ |  |  |

**Table B2. PCR Profile used for Genotyping**

| **Step** | **Temperature** | **Duration** |
| --- | --- | --- |
| Initial denaturation | 94 °C | 3 min |
| 35 cycles |  |  |
| Denaturation | 94 °C | 1 min |
| Annealing | (Optimized as per **Table 2**) | 1 min |
| Elongation | 72 °C | 1 min |
| Final elongation | 72 °C | 5 min |
